# Identification of Leucinostatins from *Ophiocordyceps* sp. as Antiparasitic Agents Against *Trypanosoma cruzi*

**DOI:** 10.1101/2021.05.30.446362

**Authors:** Jean A. Bernatchez, Yun-Seo Kil, Elany Barbosa da Silva, Diane Thomas, Laura-Isobel McCall, Karen L. Wendt, Julia M. Souza, Jasmin Ackermann, James H. McKerrow, Robert H. Cichewicz, Jair L. Siqueira-Neto

## Abstract

Safe and effective treatments for Chagas disease, a potentially fatal parasitic infection associated with cardiac and gastrointestinal pathology and caused by the kinetoplastid parasite *Trypanosoma cruzi*, have yet to be developed. Benznidazole and nifurtimox, which are currently the only available drugs against *T. cruzi*, are associated with severe adverse effects and questionable efficacy in the late stage of the disease. Natural products have proven to be a rich source of new chemotypes for other infectious agents. We utilized a microscopy-based high-throughput phenotypic screen to identify inhibitors of *T. cruzi* from a library of natural product samples obtained from fungi procured through a Citizen Science Soil Collection Program (https://whatsinyourbackyard.org/), and the Great Lakes (USA) benthic environment. We identified five leucinostatins (A, B, F, NPDG C and NPDG D) as potent inhibitors of the intracellular amastigote form of *T. cruzi*. Leucinostatin B also showed *in vivo* efficacy in a mouse model of Chagas disease. Given prior reports that leucinostatins A and B have antiparasitic activity against the related kinetoplastid *T. brucei*, our findings suggest a potential cross-trypanocidal compound class and provide a platform for further chemical derivatization of a potent chemical scaffold against *T. cruzi*.

## INTRODUCTION

Chagas disease, caused by the protozoan parasite *Trypanosoma cruzi*, remains a serious global health burden, with six to seven million individuals infected worldwide; an estimated 300,000 to 1 million cases are found in the United States alone ^1^. *T. cruzi* is usually transmitted via a blood meal by triatomine insects (commonly known as kissing bugs), but can also be acquired congenitally, through blood transfusions, or orally from ingestion of *T. cruzi-*contaminated food ^2^. Within the mammalian host, *T. cruzi* can be found in two lifecycle stages: the circulating trypomastigote stage and the intracellular amastigote stage. Parasites can persist at low levels in multiple organs including the heart and gastrointestinal tract, leading to cardiac and gastrointestinal pathology in chronically-infected patients, which can be fatal ^3,4^. Benznidazole and nifurtimox, the sole available drugs for *T. cruzi* treatment, are known to cause severe adverse effects that complicate their clinical use, and their utility in the chronic phase of infection is debated within the scientific community ^5,6^. Chagas disease is classified as a neglected tropical disease (NTD), and its historical prevalence in resource-limited countries in Latin America has restricted industrial interest in the development of a therapeutic ^1^. Given the paucity of safe and effective pharmaceutical options for *T. cruzi*, there is an urgent need for the discovery and advancement of new chemical entities for the treatment of this disease.

Natural products represent a rich source of chemical space which has yielded many successful drugs ^7^. Recent work has utilized natural product scaffolds to develop drug leads for trypanosomes. Plant and marine-derived molecules against *T. cruzi* and *T. brucei* have been successfully isolated and characterized in ongoing drug discovery ventures including derivatives of flavanones, gallinamides and polyene macrolactams ^8–10^. Furthermore, amphotericin B and paromomycin are compounds derived from bacterial sources that are used in the treatment of *Leishmania* infections ^11^.

Fungi are also an important source of antiparasitic drug scaffolds: our recent work screening for active compounds against *Trichomonas vaginalis* revealed anthraquinones, xanthone-anthraquinone heterodimers, and decalin-linked tetramic-acid-containing metabolites as moderate to potent inhibitors of this pathogen ^12^. In light of this, we screened a library of fungal natural products against *T. cruzi* to identify new chemical entities with antitrypanosomal activity. The fungal natural products were obtained via a citizen-science soil-sampling initiative (https://whatsinyourbackyard.org/), as well as a survey of Great Lakes sediments ^13,14^. Here, we report leucinostatins A, B and F, as well as novel leucinostatins NPDG C and NPDG D, which we recently characterized ^15^, as having potent activity against the replicative, intracellular amastigote form of the parasite and no host cell toxicity up to 1.5 µM using a phenotypic, high-content imaging assay. We further demonstrated that leucinostatin B prevented parasite growth in an *in vivo* bioluminescent model of *T. cruzi* infection. These results represent a framework for further development of leucinostatins and derivatives for Chagas disease treatment.

## RESULTS

### Fungal natural products library

The natural products used for bioactivity screening were derived from fungi collected using two distinct sampling strategies ^13,14^. The University of Oklahoma Citizen Science Soil Collection is a program aimed at offering citizen scientists throughout the United States the opportunity to share soil samples from which fungal isolates are obtained. Today, the collection contains fungi collected from across the country, representing every major ecological biome throughout the region. A map of sample sites and descriptions of the samples have been archived and can be viewed at the SHAREOK data sharing portal (https://shareok.org/handle/11244/28096). The University of Oklahoma natural product screening collection contains additional sets of fungi and their secondary metabolites obtained through special focused collection initiatives. One such subset was prepared from fungi derived from sediment samples collected throughout Lakes Michigan and Lake Superior, as well as a scattering of samples collected in northern Lake Huron. The natural product collection contains >78,000 samples consisting of mixtures of fungal biosynthetic products that had been prepared by subjecting cultures of the isolates to organic solvent extraction followed by partitioning using ethyl acetate-water.

### Primary screening of fungal extract samples

A subset of 5,631 samples from the University of Oklahoma fungal natural products library were tested at 2 µg/mL against *T. cruzi* strain CA-I/72 in a phenotypic high-content imaging assay we had previously developed and implemented in other drug discovery initiatives ^16–19^. Dimethyl sulfoxide (DMSO) vehicle-treated infected mouse myocytes were included as positive infection controls, while infected mouse myocytes treated with 50 µM benznidazole were included as the negative infection control. The results from this screen are presented in Figure 1; hit cutoff conditions were set at >75% antiparasitic activity and >50% host mouse myoblast cell viability. From this initial screen, a subset of 259 bioactive samples were retained for follow-up.

**Figure 1.**
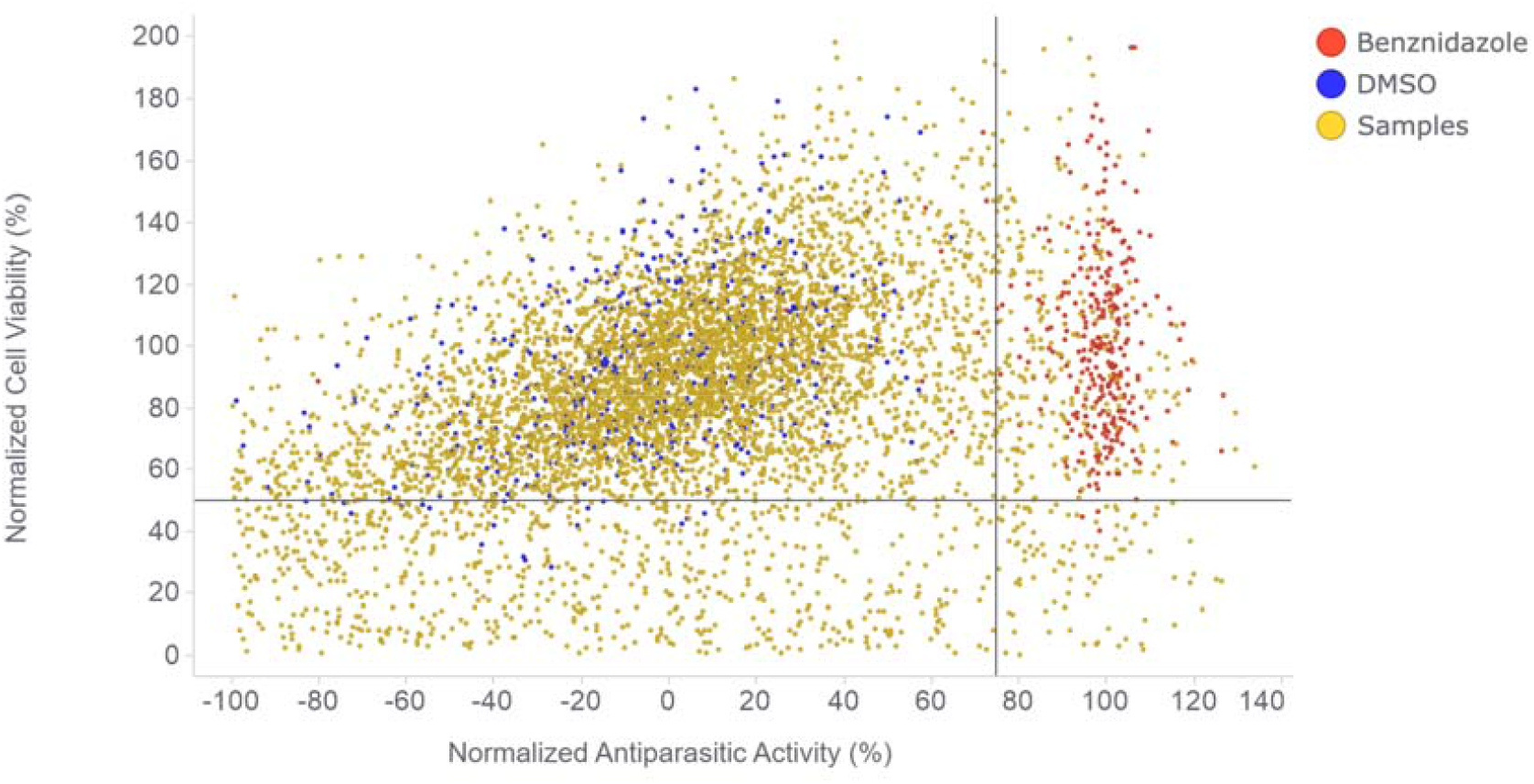
Primary screen of a fungal natural products library against *T. cruzi*. A collection of 5,631 samples from the University of Oklahoma natural product collection were screened at a final assay concentration of 2 µg/mL against *T. cruzi* CA-I/72 amastigotes using a phenotypic high-content imaging assay. Benznidazole-treated (50 µM), DMSO vehicle-treated (0.1% final concentration), and natural-product-treated samples are shown in red, blue and yellow, respectively. DMSO control-normalized percent cell viability of host cells and percent antiparasitic activity are represented on the y-axis and x-axis of the graph, respectively. Vertical and horizontal lines on the graph represent the hit threshold cutoffs for antiparasitic activity (75%) and host cell viability (50%).

### Identification of five leucinostatins from an *Ophiocordyceps* sp. as the active, pure compounds against *T. cruzi*

An iterative bioactivity-guided fraction process led to the identification of an active fraction enriched in putative peptidic natural products. Subjecting this sample to further chromatographic steps led to purification of five natural products, which included three previously reported metabolites (leucinostatins A, B and F) and two leucinostatin analogues (leucinostatins NPDG C and NPDG D) that were recently reported as new natural products ^15^ (Figure 2). The structures of the metabolites were determined using a combination of spectrometric, spectroscopic, and chemical methods ^15^. The leucinostatins and the positive control benznidazole were tested to assess their concentration-dependent antiparasitic and cytotoxic activities (Figures 3 and 4). All five leucinostatins exhibited EC_50_ values in the low nanomolar range, with no host cell toxicity detected at concentrations up to 1.5 µM, which translated into a good selectivity index of >120 for all hits (Table 1).

**Figure 2.**
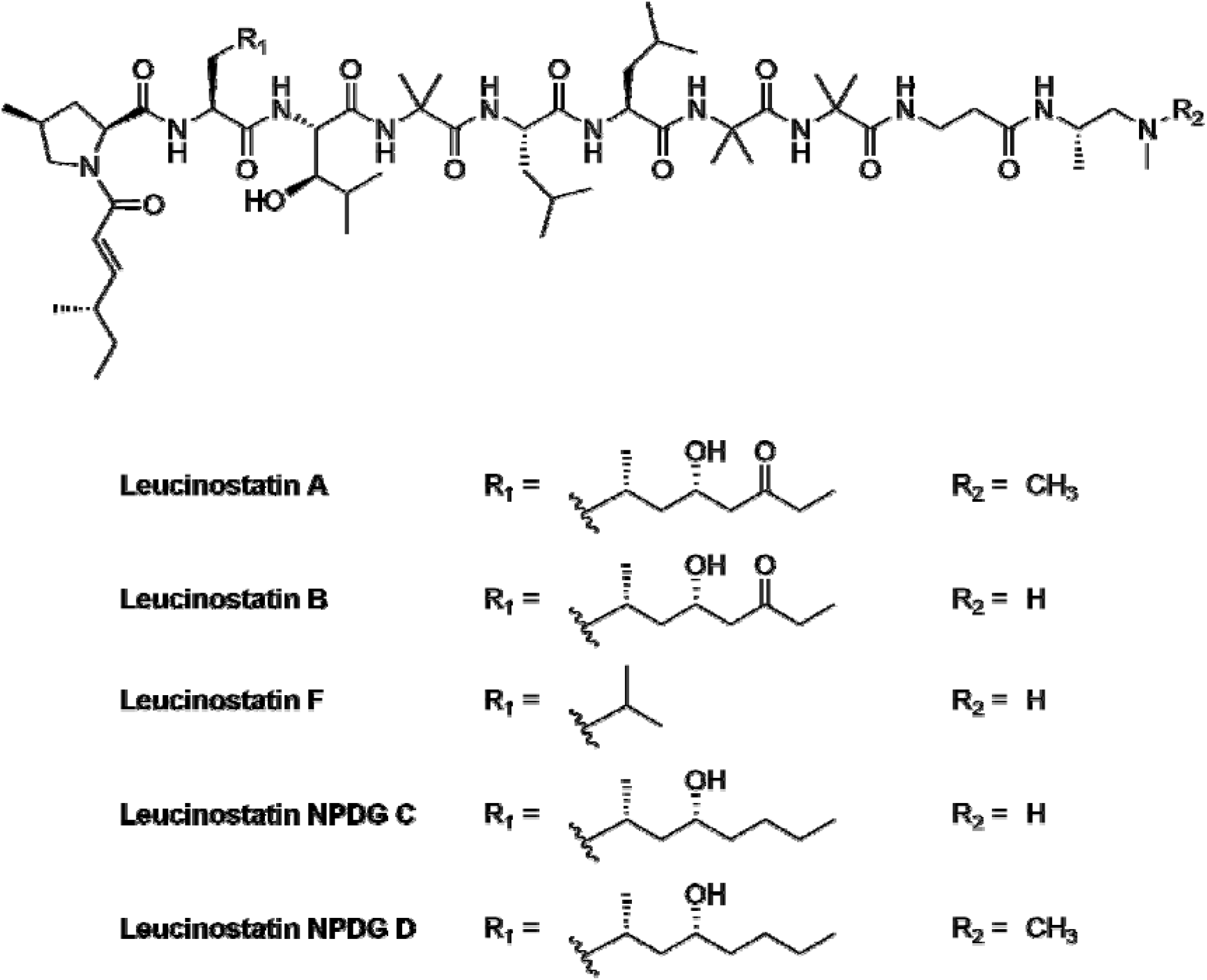
Chemical structures of the five leucinostatins active against *T. cruzi*.

**Figure 3.**
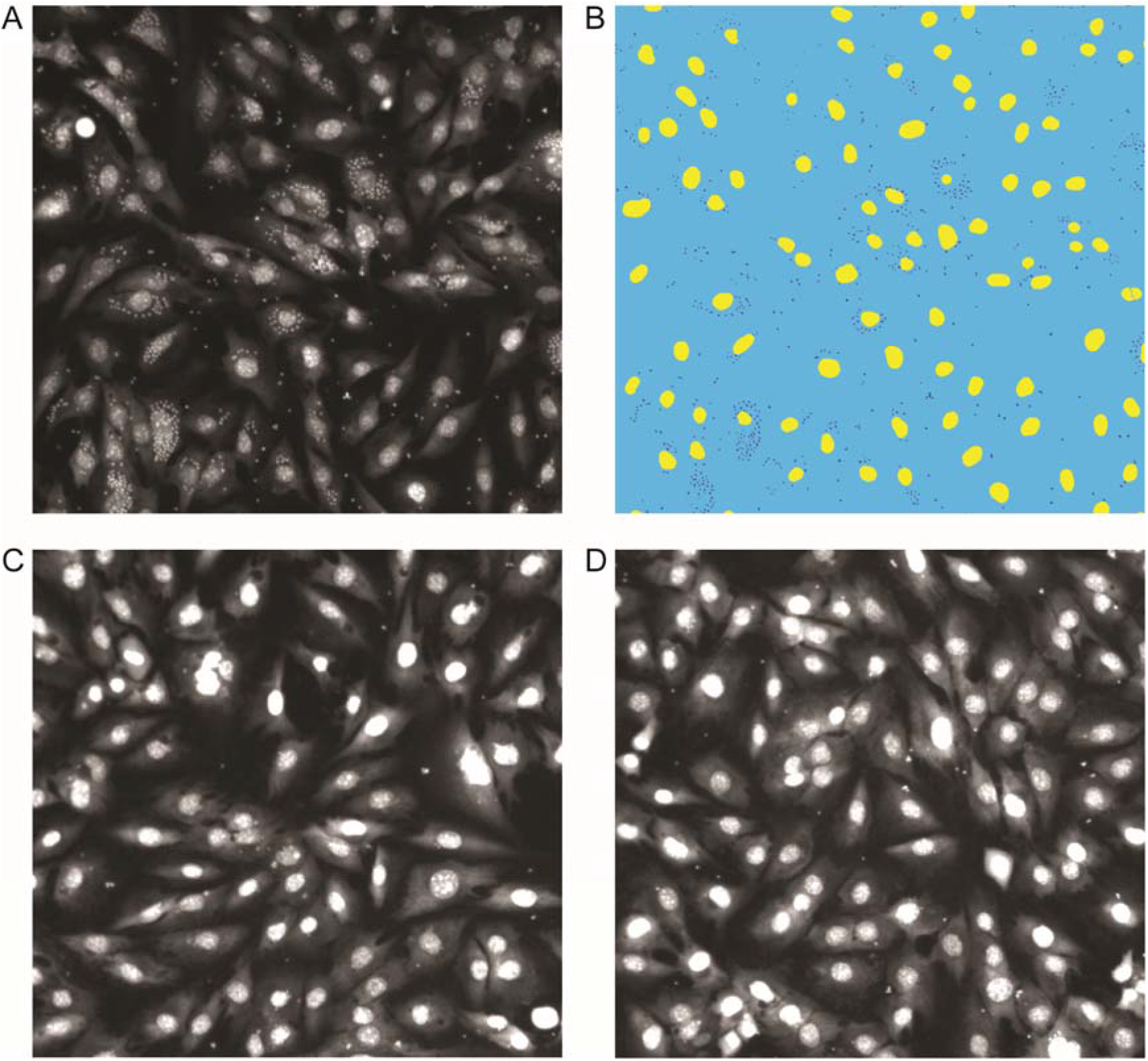
Leucinostatins inhibit *T. cruzi* replication *in vitro*. DAPI-stained host cells and parasites were imaged using an ImageXpress Micro XLS automated microscope at 10x magnification. A. DMSO vehicle-treated C2C12 myocytes infected with CA-I/72 *T. cruzi*. B. Mask of the custom automated image analysis module used to count the number of host cells and intracellular *T. cruzi* amastigotes in each microscopy image. Host cell nuclei are colored in yellow and *T. cruzi* nuclei are colored in dark blue. C. Benznidazole-treated (50 µM) C2C12 cells infected with *T. cruzi*. D. Leucinostatin F-treated (18 nM) C2C12 cells infected with *T. cruzi*.

**Figure 4.**
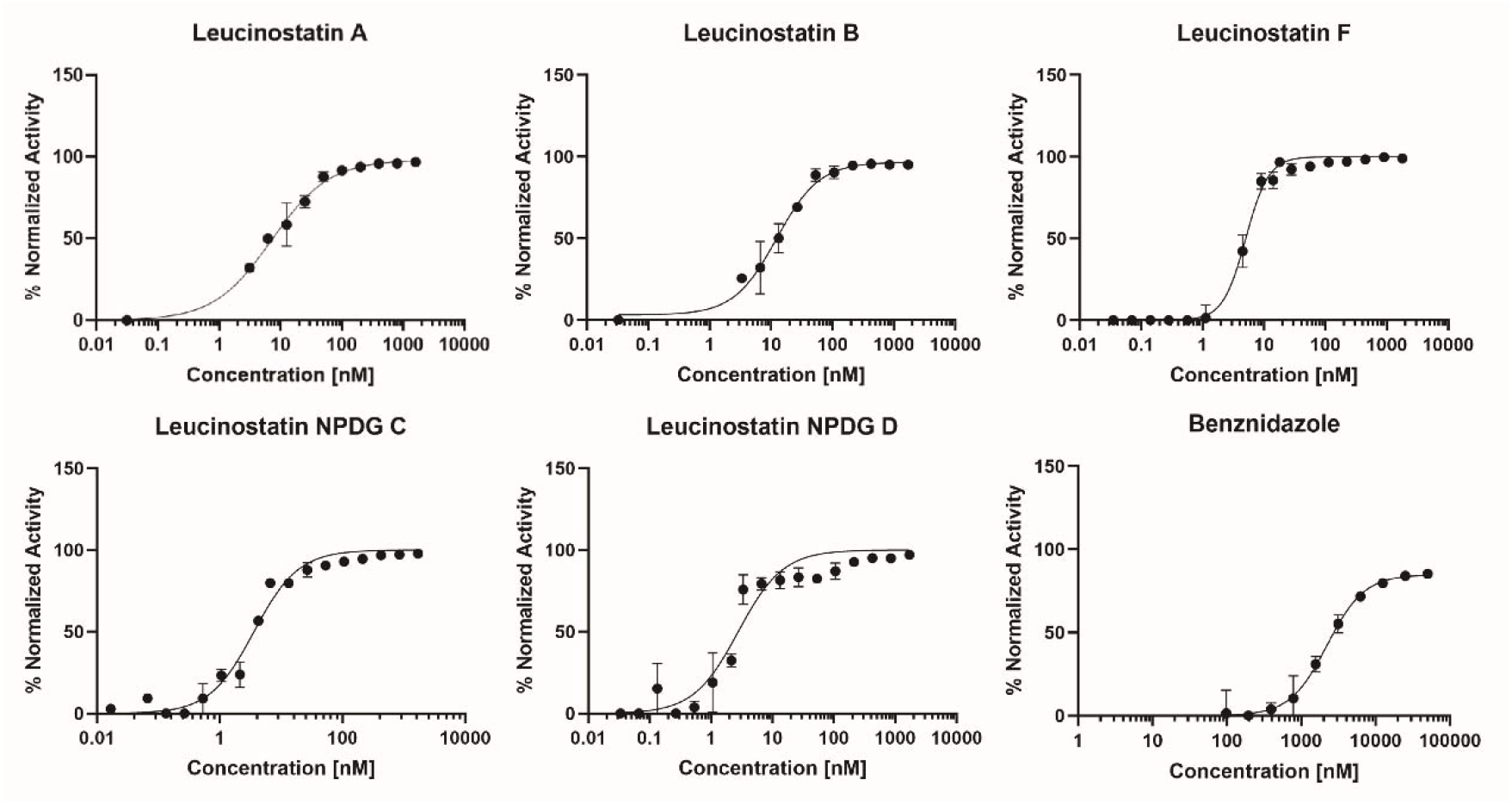
Dose-response curves for the antiparasitic activity of leucinostatin hits identified from the high-throughput screening campaign and benznidazole. Experiments were performed in 2-4 biological replicates, with bars representing the standard error of the mean for each data point shown on the graphs.

**Table 1.** Inhibition of *T. cruzi* amastigote replication by leucinostatins A, B, F, NPDG C, NPDG D and benznidazole. EC_50_ and CC_50_ are the average of two independent biological replicates, ± standard error of the mean (SEM). Selectivity index (SI) = CC_50_/EC_50_.

| Compounds | EC<sub>50</sub> (nM) | CC<sub>50</sub> (nM) | SI |
| --- | --- | --- | --- |
| **Leucinostatin A** | 7.1 ± 1.6 | >1500 | >210 |
| **Leucinostatin B** | 12 ± 1.4 | >1500 | >120 |
| **Leucinostatin F** | 5.0 ± 1.1 | > 1500 | >300 |
| **Leucinostatin NPDG C** | 3.6 ± 1.2 | > 1500 | >410 |
| **Leucinostatin NPDG D** | 2.8 ± 1.4 | > 1500 | >530 |
| **Benznidazole** | 2200 ± 1.3 | >50000 | >22 |

### *In vivo* efficacy proof-of-concept study

To test if the *in vitro* antiparasitic activity of the leucinostatins would translate to *in vivo* efficacy, we used a mouse model of acute Chagas disease ^20^. Mice were infected with luciferase-expressing *T. cruzi* trypomastigotes, sorted into groups to equalize the parasite load three days post infection, and dosed by i.p. injection with either vehicle, benznidazole, or leucinostatin B from days 3-6 post infection. The doses of leucinostatin B chosen for the *in vivo* efficacy study were based on previously published experiments that tested the efficacy of leucinostatin B in doses ranging from 0.3 mg/kg up to 2.5 mg/kg by i.p. administration for up to four days against *T. brucei* infection in mice ^21^. Therefore, we employed a dosing regimen with escalating doses starting at 0.25 mg/kg at day 3 post infection, increasing to 0.5 mg/kg at day 4 post infection, and topping out at 1 mg/kg at days 5 and 6 post infection to minimize the risk of toxicity while maximizing the chance of reaching a therapeutic dose. Leucinostatin B successfully reduced parasite growth compared to vehicle-treated mice (Figure 5). A significant difference in the slope of parasite-derived luminescence over time was observed between benznidazole and vehicle groups (p=0.0007) and between leucinostatin B and vehicle groups (p=0.02). No significant difference in slope was observed for benznidazole to leucinostatin B groups (p=0.6).

**Figure 5.**
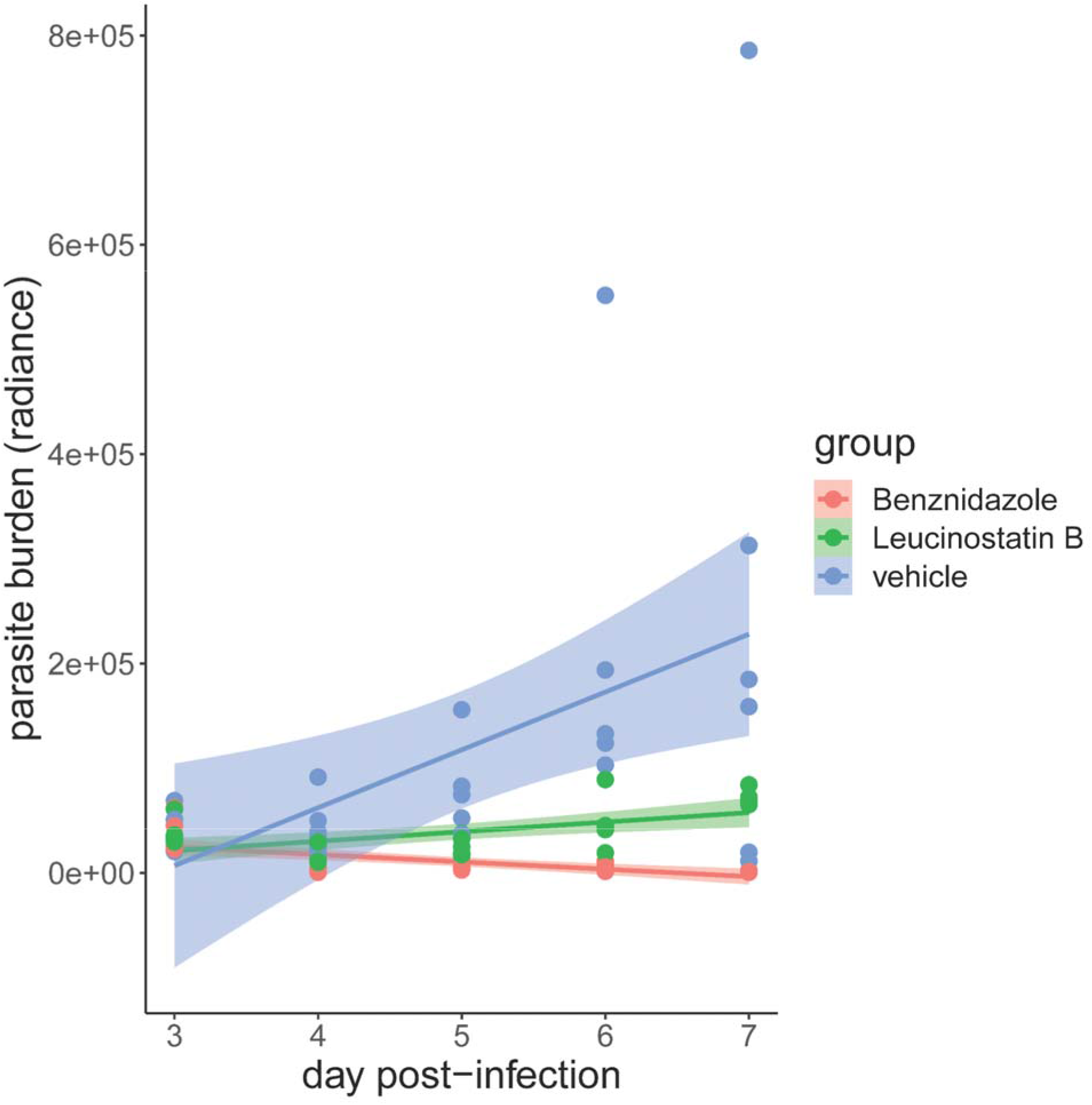
Antiparasitic efficacy of leucinostatin B in a mouse model of Chagas disease. *T. cruzi* infected mice were treated beginning 3 days post-infection with benznidazole (50 mg/kg b.i.d., i.p.), leucinostatin B (escalating dose regimen: 3 dpi: 0.25 mg/kg; 4 dpi: 0.5 mg/kg; 5 and 6 dpi: 1 mg/kg; all b.i.d., i.p.), or vehicle (10% DMSO, b.i.d. i.p.). Parasite load was quantified by bioluminescence daily. Shaded areas represent 95% confidence intervals, all sexes combined.

## DISCUSSION

Treatment options for Chagas disease remain limited. Efforts to expand the scope of chemical scaffolds for drug discovery programs against *T. cruzi* using natural product, *de novo* design, and drug repositioning approaches have yielded promising new leads in recent years ^9,16,22–25^. Here, we used a high-throughput screening and bioactivity-guided fractionation pipeline to identify five potent leucinostatins from an *Ophiocordyceps* sp. with low nanomolar EC_50_ values and no host cell toxicity at concentration up to 1.5 µM. These EC_50_ values are particularly promising, since selection criteria for *T. cruzi* screening campaigns usually consider <10 µM EC_50_ values as hit molecules ^26^. Benznidazole EC_50_ of 2.2 µM as a point of comparison demonstrates how much more potent the leucinostatins are as *T. cruzi* inhibitors than the reference drug. Furthermore, the large selectivity index ranging from >120 to >530 for the leucinostatin hits in our study greatly exceeded standard hit selection criteria for emerging infectious diseases (SI of >10) ^26^. Using a mouse model of Chagas disease, we were also able to demonstrate significant antiparasitic activity *in vivo*, which is an encouraging proof-of-concept for future development of this chemical scaffold as a therapeutic agent.

Previous work has shown that both leucinostatins A and B have activity against the causative agent of African trypanosomiasis, *T. brucei* ^*21*^; the authors of the study speculated that the mechanism of action of these compounds may involve targeting of the ATP synthetase and/or Ca^2+^ and pH homeostasis in *T. brucei*. We have confirmed that both these compounds inhibit the related kinetoplastid *T. cruzi*, and discovered three additional leucinostatin scaffolds. These findings suggest that a potential common mechanism of inhibition across trypanosomes for this class of compounds. Future studies will focus on testing these compounds for potential activity against the related kinetoplastid *Leishmania* sp. and mechanism of action studies. In addition, SAR studies and pharmacokinetic optimization will be logical next steps in the development of these compounds as potential anti-Chagas drug leads while ensuring adequate drug tolerability.

In summary, we have determined that naturally-occurring leucinostatins are promising natural products with potent *in vitro* and *in vivo* activities against *T. cruzi*, and cross-trypanosome activity. These results set the stage for the development of this class of antitrypanosomal molecules as leads for the treatment of Chagas disease.

## EXPERIMENTAL SECTION

### Cells

CA-I/72 *T. cruzi* (a gift from J. Dvorak, NIH) and C2C12 mouse myoblasts (ATCC CRL-1772) were passaged in Dulbecco’s Modified Eagle Medium (Invitrogen, 11095-080) with 5% HyClone iron-supplemented calf serum (Cytivia, SH30073.03) and 1% penicillin-streptomycin (Invitrogen, 15140122), and were maintained at 37 °C and 5% CO_2_. CA-I/72 *T. cruzi* were passaged weekly through co-culture with C2C12 mouse myoblasts, essentially as described ^20^.

### Fungi and natural products

The methods related to the identification and culture of the *Ophiocordyceps* sp., purification of the leucinostatins, and methods used to solve the structures of the natural products have been reported ^15^.

### *In vitro* antiparasitic assay

Samples and pure compounds from the fungal natural product library, benznidazole (Sigma cat. no. 419656) and DMSO (Sigma cat. no. D2650) were spotted onto black, clear bottom 384-well plates (Greiner Bio One, 782092) using an Acoustic Transfer System (ATS) instrument (EDC Biosystems). Using a Multidrop Combi liquid handler (Thermo Scientific), 500 cells per well of C2C12 cardiomyoblasts and 7,500 cells per well of CA-I/72 *T. cruzi* parasites were added to each well plate. This was followed by incubation at 37 °C and 5% CO_2_ for 72 hours, using plate-holding trays to reduce evaporation of cultures at the edges of the plates. Paraformaldehyde (4% final concentration) in 1x phosphate buffered saline (PBS, Invitrogen, 10010023) was then added to the plates to fix the cells. Subsequent treatment with 5 µg/mL DAPI staining solution (Sigma Aldrich, D9542) was applied for 1 hour to stain host cell and parasite nuclei. Imaging of well plates was conducted with a 10x fluorescence objective using an ImageXpress Micro XLS automated high-content imager (Molecular Devices). A custom image analysis module generated in MetaXpress (Molecular Devices) was then used to count host cell and parasite nuclei to be used as an assay readout, as previously described ^20^.

### *In vivo* antiparasitic proof-of-concept study

Six-week-old BALB/c mice (3 males and 3 females per group) were each infected with 1 × 10^5^ trypomastigotes of PpyRE9h luciferase-expressing *T*.□*cruzi* CL Brener ^27^. Three days post infection (dpi) the mice were sorted to equalize parasite load into 3 groups and received the following treatments for 4 consecutive days (from 3 dpi to 6 dpi): vehicle, 10% DMSO, bis-in-die (b.i.d.), intraperitoneally (i.p.); benznidazole, 50 mg/kg, b.i.d., i.p.; and leucinostatin B in an escalating dose regimen: 3 dpi: 0.25 mg/kg; 4 dpi: 0.5 mg/kg; 5 and 6 dpi: 1 mg/kg; all b.i.d., i.p.). Mice were each injected (i.p.) with 100 µl of 25 mg/ml D-luciferin potassium salt (GoldBio, St. Louis, MO) and imaged daily from days 3 dpi - 7 dpi with an In Vivo Imaging System (IVIS, PerkinElmer) to measure luminescence signal corresponding to parasite burden. The change in average radiance over time for each treatment group was analyzed by linear regression analysis and slopes compared using the ‘lsmeans’ package in R. p-Value adjustment was performed using the Tukey method. The plot was generated using the ‘ggplot’ package in R.

### Software

GraphPad Prism 8 (GraphPad Software) was used to generate EC_50_ and CC_50_ curves for the phenotypic data. CDD Vault (Collaborative Drug Discovery Inc.) was used to curate primary screening and counter-screening data. *In vivo* data was analyzed using R version 3.6.1 in Jupyter Notebook.

## Author Contributions

^¶^J.A.B. and Y.-S.K. contributed equally. All authors wrote and revised the manuscript.

## Notes

The authors declare no competing financial interest.

### Ethics Statement

All animal studies were performed under approved protocol S14187 from the Institutional Animal Care and Use Committee, University of California, San Diego (AAALAC Accreditation Number 000503) and in compliance with the Animal Welfare Act and adhere to the principles stated in the Guide for the Care and Use of Laboratory Animals, National Research Council, 2011.

## Acknowledgments

We thank Dr. Lars Eckmann from the University of California, San Diego, CA, USA for the gift of the BALB/c mice used in the *in vivo* studies. We thank Dr. John Kelly from the London School of Hygiene and Tropical Medicine, London, UK for the gift of the PpyRE9h luciferase-expressing *T*.□*cruzi* CL Brener.

